# Pre-existing immunity modulates responses to mRNA boosters

**DOI:** 10.1101/2022.06.27.497248

**Authors:** Tanushree Dangi, Sarah Sanchez, Min Han Lew, Lavanya Visvabharathy, Justin Richner, Igor J. Koralnik, Pablo Penaloza-MacMaster

## Abstract

mRNA vaccines have shown high efficacy in preventing severe COVID-19, but breakthrough infections, emerging variants and waning antibody levels have warranted the use of boosters. Although mRNA boosters have been widely implemented, the extent to which pre-existing immunity influences the efficacy of boosters remains unclear. In a cohort of individuals primed with the mRNA-1273 or BNT162b2 vaccines, we observed that lower antibody levels before boost were associated with higher fold-increase in antibody levels after boost, suggesting that pre-existing antibody modulates the boosting capacity of mRNA vaccines. Mechanistic studies in mice show that pre-existing antibodies significantly limit antigen expression and priming of B cell responses after mRNA vaccination. Furthermore, we demonstrate that the relative superiority of an updated Omicron vaccine over the original vaccine is critically dependent on the serostatus of the host. These data demonstrate that pre-existing immunity dictates responses to mRNA vaccination, elucidating specific circumstances when updated SARS-CoV-2 vaccines confer superior protection to original vaccines.

## Introduction

mRNA lipid nanoparticle (RNA-LNP) vaccines have been administered to millions of people worldwide, showing high efficacy against COVID-19. The mRNA-LNP platform has revolutionized multiple fields of medicine, including vaccinology, cancer therapy and gene therapy. Despite their wide use, the immunobiology of mRNA-LNPs remains incompletely understood, especially regarding how pre-existing immunity elicited by prior vaccination or infection can affect the efficacy of mRNA vaccines or responses to updated boosters. This knowledge would be critical during the next phase of the COVID-19 pandemic, as vaccine manufacturers are currently testing updated mRNA boosters based on variant sequences to determine whether they can confer an immunological advantage over the ancestral vaccines.

Both Pfizer-BioNTech and Moderna have started vaccine trials to evaluate Omicron-based vaccines for the prevention of Omicron infection. Moderna has recently released preliminary data on its Phase 2/3 trial (NCT05249829), which suggested that an updated bivalent booster based on both Omicron and ancestral spike antigens elicits superior neutralizing antibody against Omicron than the ancestral vaccine. However, other studies have suggested that when given as a third shot, Omicron-based vaccines may not necessarily confer superior protection to the original vaccine (*1–3*). Here, we aimed to answer two critical questions that are important in the current phase of the COVID-19 pandemic, while Omicron-based vaccines seek licensure. First, how does pre-existing immunity affect responses to mRNA vaccines? Second, are there specific circumstances where updated vaccines are more effective than ancestral vaccines? We show that that pre-existing immunity can impinge upon the efficacy of mRNA vaccines, and that Omicron vaccines can confer an immunological advantage in seronegative hosts. These data may provide important insights for improving the efficacy of mRNA vaccines.

## Main points

1. **In human volunteers who received Pfizer-BioNTech or Moderna vaccines, antibody levels before boost are inversely correlated to their fold-increase after boost.**
2. **A similar inverse association was observed in COVID-19 convalescent individuals who then received Pfizer-BioNTech or Moderna vaccines.**
3. **Pre-existing antibody limits antigen expression and de novo B cell responses following mRNA vaccination.**
4. **Omicron vaccines confer superior protection against Omicron relative to ancestral vaccines, when administered in a seronegative host.**

## Results

### Low pre-boost antibody levels are associated with greater fold-increase in antibody levels post-boost

Despite effective vaccines, SARS-CoV-2 continues to spread and mutate around the world. This has motivated additional boosters, but little is known about how pre-existing immunity affects responses elicited by boosters. We first interrogated whether the level of pre-existing immunity to the SARS-CoV-2 spike antigen would affect the boosting capacity of mRNA vaccines in a cohort of unexposed (COVID-19 negative) individuals who previously received one dose of mRNA vaccine (Figure 1A). Interestingly, volunteers who exhibited the lowest spike-specific antibody response before boost showed the highest fold increase in spike-specific antibody after boost (Figure 1B).

**Figure 1.**
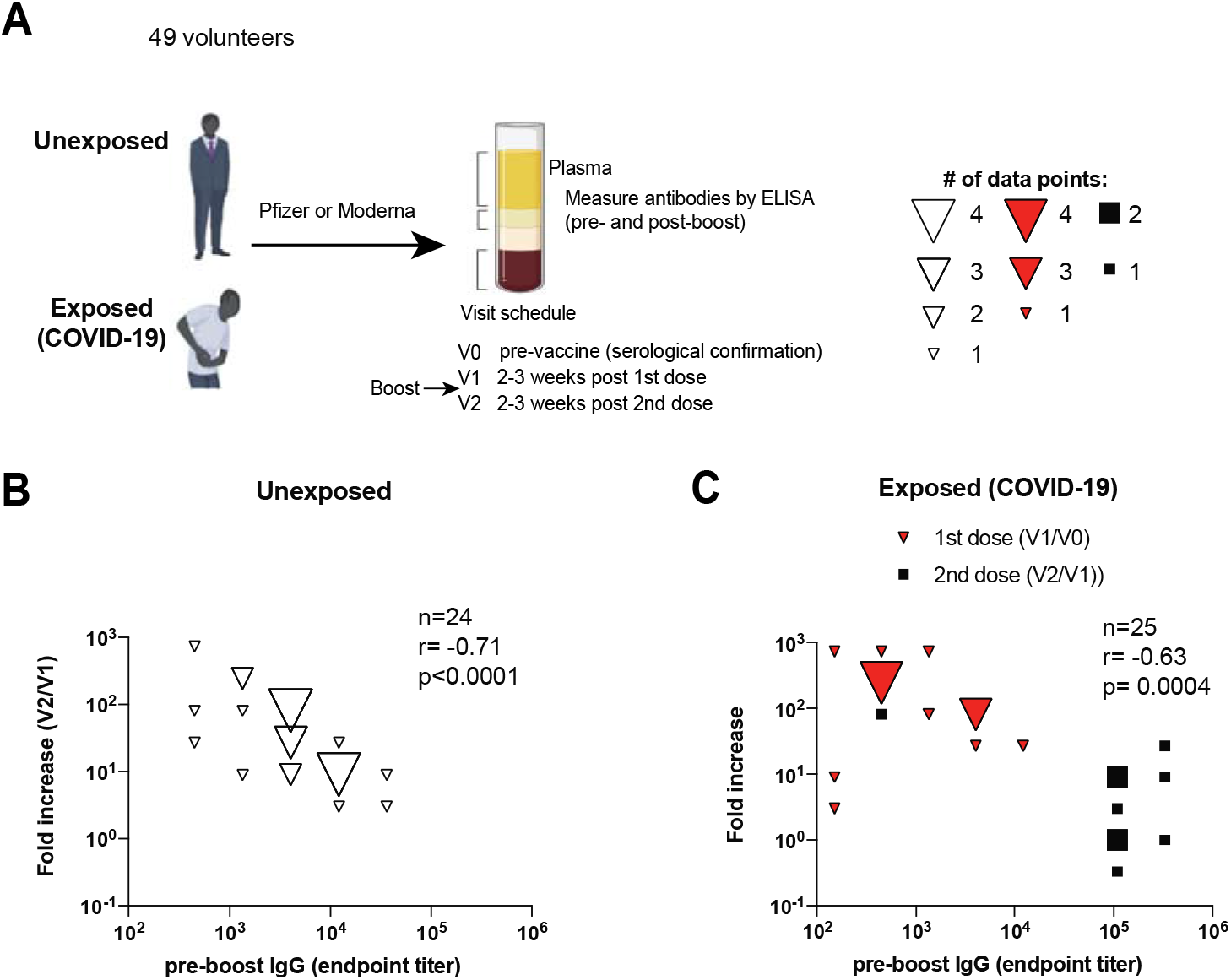
Pre-boost antibody levels are inversely correlated to fold-increase in antibody levels following mRNA vaccination in humans. (A) Experimental layout. Participants were determined to be unexposed prior to vaccination based on a negative serology test for SARS-CoV-2 spike and nucleocapsid proteins before vaccination. Participants were determined to be exposed to SARS-CoV-2 based on symptoms and confirmatory RT-PCR. (B) Summary of SARS-CoV-2 spike antibody responses in unexposed individuals. (C) Summary of SARS-CoV-2 spike antibody responses in exposed individuals. Since endpoint titers fall on discrete values (multiples of 25), several values overlapped on the same data point, so bubble plots were utilized to depict overlapping data points. Fold increase was calculated by dividing the post-boost antibody titer by the pre-boost antibody titer (SARS-CoV-2-spike specific IgG). Data are from 3 visits or time points (V0, V1, V2). Non-parametric Spearman correlation was used to calculate correlation between pre-boost antibody titer and fold-increase in antibody titers (two-tailed test was used to calculate significance).

We also analyzed a cohort of individuals with prior infection with SARS-CoV-2, and we observed that patients who exhibited the lowest antibody response following infection (pre-vaccination), showed the highest fold-increase in antibody response after a first dose of vaccine (Figure 1C). A similar inverse association was observed in SARS-CoV-2 infected individuals who received a second dose of vaccine (Figure 1C). Collectively, these data demonstrate an inverse association between pre-existing antibody levels and the ability of mRNA vaccines to boost antibody responses. A possible reason for such negative correlation could be that as antibody responses reach a certain threshold, it could be increasingly more difficult for the antibody response to undergo further recall expansion. Another reason could be that pre-existing antibodies may accelerate the clearance of vaccine antigen, limiting the access of B cells to their antigen. To better understand how seropositivity affects the immunogenicity of mRNA vaccines, we performed passive immunization studies.

### Immune plasma from mRNA-1273 or BNT162b2 vaccinated individuals abrogates *de novo* priming of B cell responses

Our data from vaccinated humans suggested that pre-existing antibody to the vaccine antigen (SARS-CoV-2 spike) can influence antibody responses following booster vaccination. To evaluate the effect of pre-existing antibody more rigorously, we transferred donor-matched human plasma (pre-vaccination plasma versus post-vaccination plasma) into naïve C57BL/6 mice; and on the following day, recipient mice received an mRNA vaccine expressing the SARS-CoV-2 spike protein (Figure 2A). Note that human- and mouse-derived antibodies can be distinguished using host-specific enzyme-linked immunosorbent assays (ELISA), allowing us to track donor-derived and recipient-derived antibodies over time.

**Figure 2.**
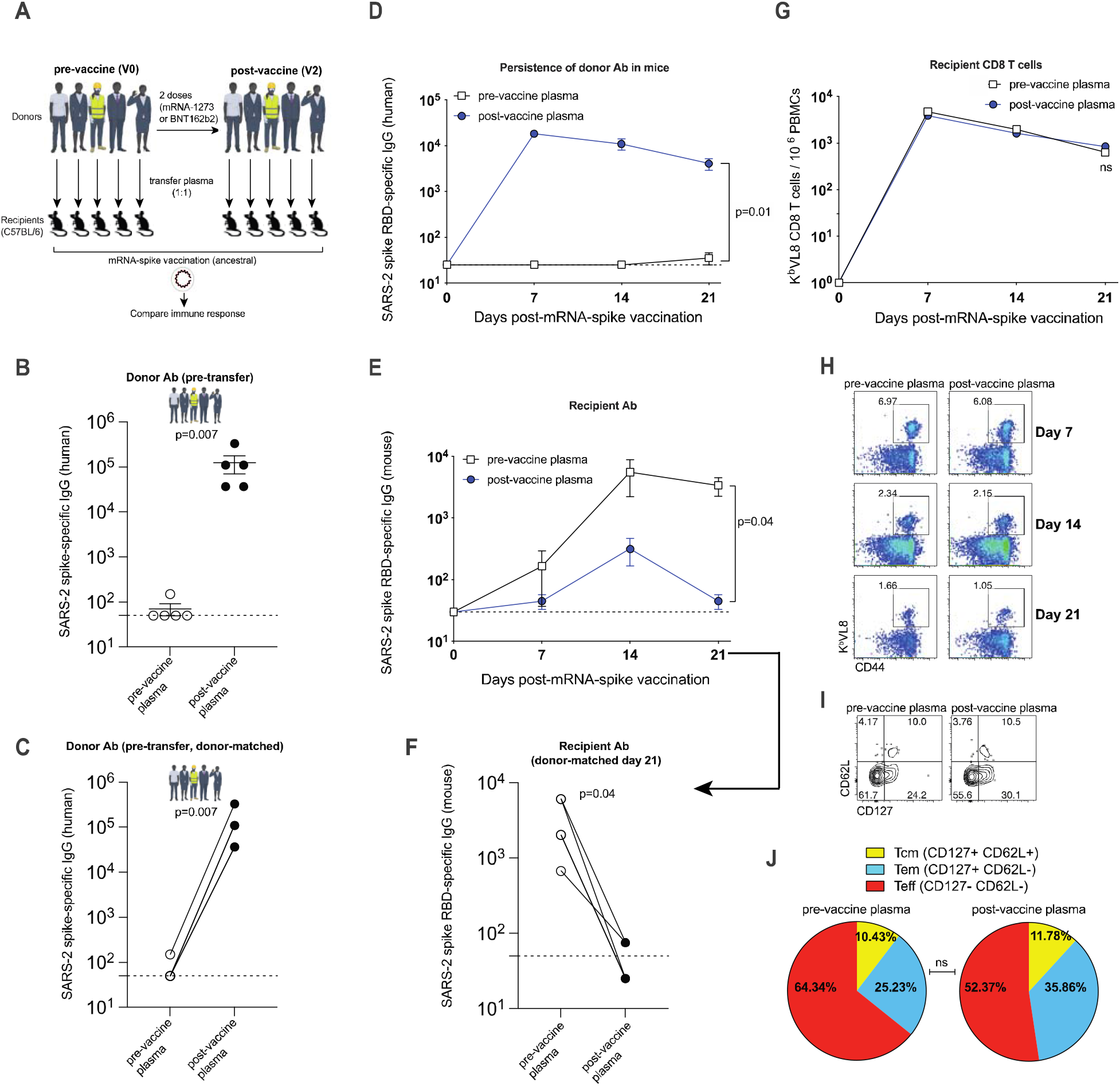
Plasma from mRNA-1273 or BNT162b2 vaccinated humans abrogates *de novo* antibody responses following mRNA vaccination in mice. (A) Experimental layout. Donor-matched plasmas were harvested from 5 different human donors; pre-vaccination (V0) and post-vaccination (V2 corresponding to 2-3 weeks post 2^nd^ dose). 100 μL of these donor-matched human plasmas were adoptively transferred via the intraperitoneal route into C57BL/6 mice. On the following day, all mice were immunized intramuscularly with 3 μg of an mRNA expressing SARS-CoV-2 spike; and immune responses were quantified every week. (B-C) SARS-CoV-2 spike–specific antibody responses in human donors who received mRNA-1273 or BNT162b2 vaccines; since endpoint titers fall on discrete values (multiples of 25), several values overlapped on the same data point, so data are shown as single replicates (B) or as donor-matched data points to visualize vaccine-elicited responses in each human donor (C). (D) Human-derived SARS-CoV-2 spike–specific antibody in mice that received plasma from mRNA-1273 or BNT162b2 vaccinated humans. (E-F) Mouse-derived SARS-CoV-2 spike– specific antibody in mice that received plasma from mRNA-1273 or BNT162b2 vaccinated humans; Data are shown as summary plots (E) or as donor-matched data points for day 21 (F). (G) Summary of SARS-CoV-2 spike–specific CD8 T cells in mice that received plasma from mRNA-1273 or BNT162b2 vaccinated humans. These CD8 T cells were specific for a CD8 T cell epitope derived from the spike protein (K^b^VL8). (H) Representative FACS plots of SARS-CoV-2 spike–specific CD8 T cells in mice that received plasma from mRNA-1273 or BNT162b2 vaccinated humans. FACS plots are gated on live CD8 T cells. (I) Representative FACS plots showing memory CD8 T cell differentiation in mice that received plasma from mRNA-1273 or BNT162b2 vaccinated humans. FACS plots are gated on live K^b^VL8+ CD8 T cells. (J) Summary of memory markers on K^b^VL8+ CD8 T cells at day 21 post-vaccination. Two-tailed parametric test (matched) was used to calculate significance in panels D-G. For all other plots, two-tailed Mann Whitney test was used. Data are from an experiment with 5 mice that received pre-vaccine human plasma and 5 mice that received post-vaccine human plasma; data from all experiments are shown, and dashed lines represent the LOD. Error bars indicate the SEM.

We first measured antibody responses in our 5 human donors, before and after mRNA vaccination to corroborate seroconversion in these individuals. As expected, we detected a robust antibody response in all 5 human donors after the second mRNA dose (Figure 2B-2C). Following transfer of human plasma into mice, we were able to detect human spike-specific antibody in all mice for up to 21 days (Figure 2D). Interestingly, mice that received pre-vaccine human plasma showed robust antibody responses after mRNA vaccination, whereas mice that received post-vaccine human plasma showed a significant impairment in antibody responses (Figure 2E-2F). These data suggested that pre-existing antibodies elicited by the mRNA-1273 or BNT162b2 vaccines can compete with *de novo* antibody responses following a booster vaccination. We did not observe any significant differences in the levels of CD8 T cells (Figure 2G-2H) or their memory differentiation (Figure 2I-2J).

Both primary and secondary B cell responses can be recruited to the immune response following a booster immunization (*4*). We utilized another experimental model that allowed us to track *de novo* B cell responses using the same donor and recipient animal species (Figure 3A). We first primed C57BL/6 mice with an mRNA-based SARS-CoV-2 vaccine expressing the ancestral spike sequence (mRNA-spike), and then boosted these at week 3. At week 2 post-boost, we harvested immune plasma from these vaccinated mice and transferred it into naïve recipient BALB/c mice. On the following day, the recipient mice were immunized with the mRNA-spike vaccine and *de novo* immune responses were measured. Note that donor-derived and recipient-derived antibodies can be distinguished based on their immunoglobulin allotype. Antibodies from donor C57BL/6 mice contain the IgG^[b]^ allele, whereas antibodies from recipient BALB/c mice contain the IgG^[a]^ allele. These two alleles can be distinguished using allotype-specific ELISAs, allowing us to track donor-derived from recipient-derived antibodies over time.

**Figure 3.**
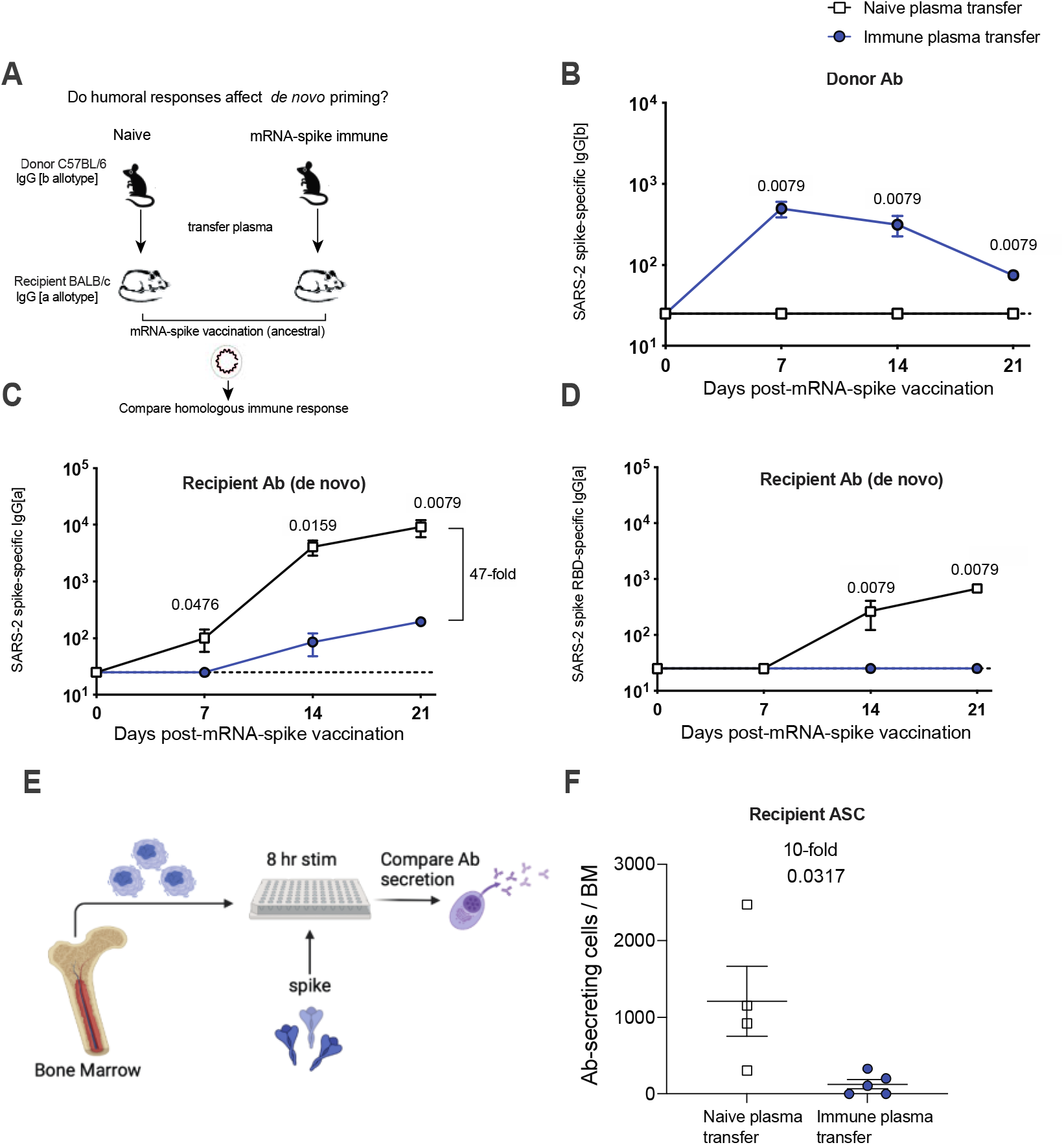
Plasma from vaccinated mice abrogates *de novo* antibody responses. (A) Experimental layout. Plasmas were harvested from C57BL/6 mice that were vaccinated with mRNA-spike (two doses). 400 μL of these plasmas were adoptively transferred via the intraperitoneal route into BALB/c mice. On the following day, all mice were immunized intramuscularly with 3 μg of an mRNA expressing SARS-CoV-2 spike; and immune responses were quantified every week. Naïve plasmas were used as control. (B) Donor-derived SARS-CoV-2 spike–specific antibody. (C-D) Recipient-derived SARS-CoV-2 spike–specific antibody; whole spike-specific (C) and RBD-specific (D). (E) Experimental layout for detection of antibody secreting cells. (F) Antibody secreting cells in bone marrow. Two-tailed Mann Whitney test was used. Data are from an experiment with 4 mice that received naïve plasma and 5 mice that received post-vaccine plasma (week 2 post-boost); experiment was repeated for a total of 2 times, with similar results; dashed lines represent the LOD. Error bars indicate the SEM.

As expected, donor antibody was detectable in recipient mice (Figure 3B). Similar to our prior experiments, mice that received naïve plasma showed robust antibody responses after mRNA vaccination, whereas mice that received immune plasma showed significantly impaired antibody responses (Figure 3C-3D) These data demonstrate that seropositivity to the vaccine antigen impairs *de novo* antibody responses following mRNA vaccination. After immunization, long-term humoral immunity is maintained by a subset of B cells that reside mostly in the bone marrow, called plasma cells (*5*). We quantified plasma cells at week 3 post post-immunization using an ELISPOT-based antibody secreting cell assay (Figure 3E), and we observed a 10-fold reduction in the number of vaccine-induced plasma cells in mice that received immune plasma (Figure 3F). We did not detect differences in T cell responses (Figure S1A-S1B)

Vaccines expressing the original spike protein of SARS-CoV-2 also generate cross-reactive Omicron-specific responses (*6–9*), and consistent with these prior reports, we observed that the ancestral vaccine elicited Omicron-specific antibody responses. However, transfer of plasma from mice vaccinated with the ancestral vaccine completely abrogated these cross-reactive antibody responses (Figure S2A-S2B) and cross-reactive plasma cell responses (Figure S2C-S2D). We detected cross-reactive Omicron-specific T cell responses in all mice, and we did not observe differences between the two groups (Figure S2E-S2F). Altogether, these data demonstrate that pre-existing humoral responses limit the priming of naïve B cell responses (but not T cell responses) following mRNA vaccination.

### Effects of pre-existing immunity on Omicron-based vaccines

As the human population reaches immunity to SARS-CoV-2 either by vaccination or infection, it becomes critical to understand the effects of pre-existing immunity on vaccine boosters. Recently, the new B.1.1.529 (Omicron) variant was identified and it is now the dominant variant worldwide. Due to its high number of mutations, Omicron can evade neutralizing antibody responses elicited by ancestral vaccines, motivating the development of Omicron-based boosters, and little is known about how pre-existing immunity generated by ancestral vaccines affects the response to updated boosters. To bring more clarity to this issue, we primed mice with an ancestral vaccine, and then we boosted them with an ancestral or an Omicron vaccine (Figure S3A). Both booster regimens elicited comparable CD8 T cell responses (Figure S3B), but we observed differences in antibody responses. As expected, boosting with the ancestral vaccine generated superior antibody responses against the ancestral virus (Figure S3C); but boosting with the Omicron vaccine did not generate superior antibody responses against Omicron (Figure S3D). Neutralizing antibody responses against the ancestral virus were 1531-fold higher in mice that received the ancestral vaccine boost, relative to mice that received the Omicron vaccine boost (Figure S3E). We also measured neutralizing antibody responses against the Omicron virus, and we observed very low responses in all mice, with many mice showing responses near the limit of detection (Figure S3F). Although the Omicron boost generated higher neutralizing antibody than an ancestral boost, the difference was only 1.4-fold (Figure S3F) Overall, in a host that has already been immunized with ancestral vaccine, boosting with an Omicron vaccine does not appear to elicit a striking improvement in the adaptive immune response, relative to boosting with an ancestral vaccine.

We also compared immune responses in mice that received homologous prime-boost with either ancestral or Omicron vaccines (Figure 4A). We did not observe differences in the number CD8 and CD4 T cells (Figure 4B-4F), but the ancestral vaccine regimen tended to generate more polyfunctional CD8 T cell responses than the Omicron vaccine, when CD8 T cells were stimulated with cognate ancestral peptide pools (p=0.03 when comparing % of triple producing cells) (Figure 4G). No statistically significant difference in polyfunctionality was observed when CD8 T cells were stimulated with Omicron peptide pools (Figure 4H). Collectively, these data showed that both vaccines elicit comparable levels of CD8 T cells and CD4 T cells.

**Figure 4.**
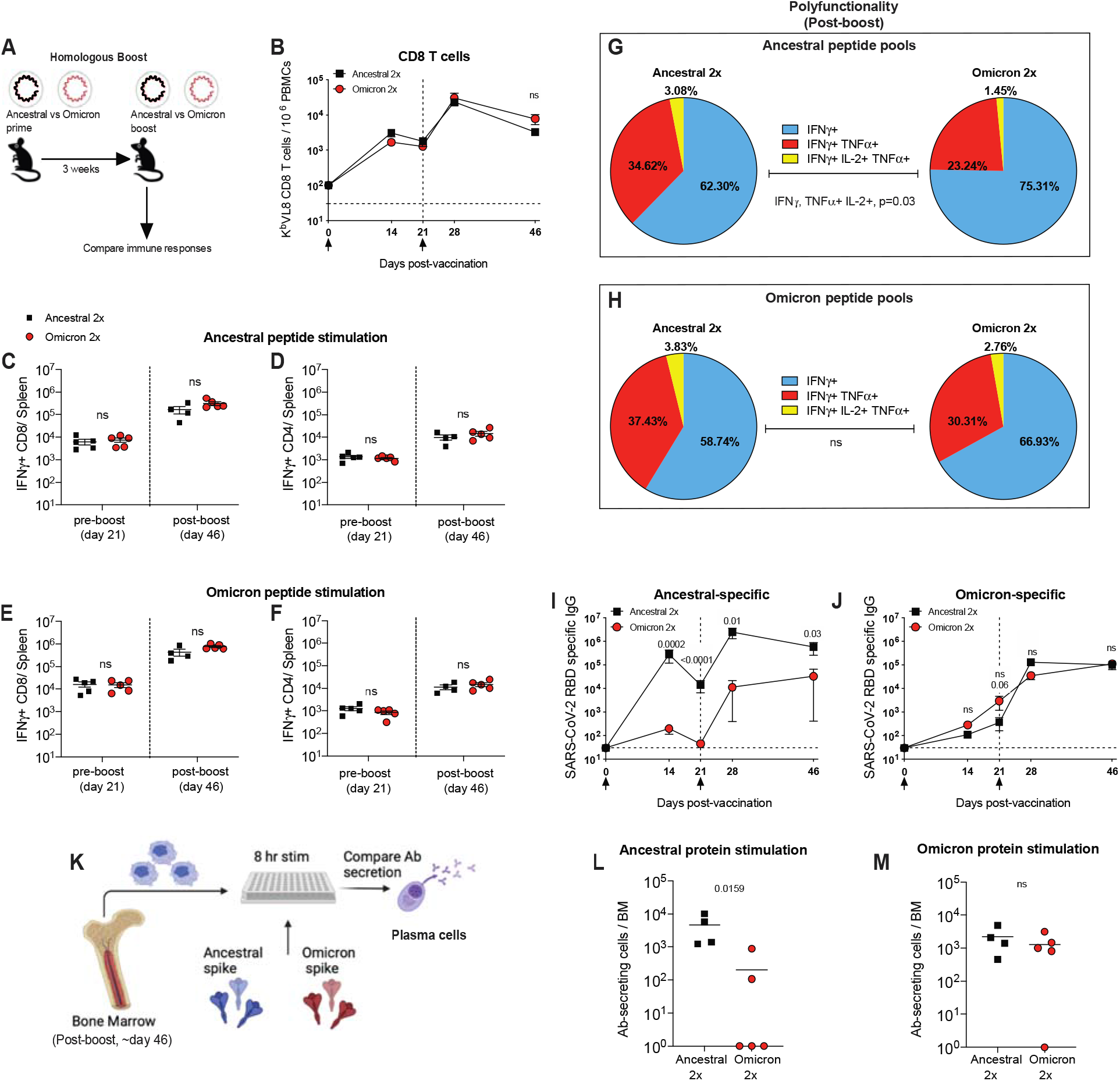
A homologous Omicron boost is not superior to a homologous ancestral vaccine boost. (A) Experimental layout. C57BL/6 mice were immunized intramuscularly with 3 μg of an mRNA expressing ancestral spike or Omicron spike. After 3 weeks, mice were boosted homologously and immune responses were quantified. (B) Summary of SARS-CoV-2 spike–specific CD8 T cells. These CD8 T cells were specific for a conserved CD8 T cell epitope present in both ancestral and Omicron viruses (K^b^VL8). (C-D) Summary of SARS-CoV-2 spike–specific CD8 and CD4 T cells after stimulation with ancestral spike peptide pools. (E-F) Summary of SARS-CoV-2 spike–specific CD8 and CD4 T cells after stimulation with Omicron spike peptide pools. (G) Polyfunctionality of CD8 T cells after stimulation with ancestral spike peptide pools. (H) Polyfunctionality of CD8 T cells after stimulation with Omicron spike peptide pools. (I) Ancestral spike–specific antibody responses. (J) Omicron spike–specific antibody responses. (K) Experimental layout for detection of antibody secreting cells specific for ancestral or Omicron spike. (L) Ancestral spike–specific antibody secreting cells. (M) Omicron spike–specific antibody secreting cells. Two-tailed Mann Whitney test was used. Data are from an experiment with 4-5 mice that received an ancestral vaccine and 5 mice that received an Omicron vaccine; experiment was repeated for a total of 2 times, with similar results; dashed lines represent the LOD. Error bars indicate the SEM.

At the end of the homologous prime-boost experiments, two shots of ancestral vaccine generated higher ancestral-specific antibody responses, relative to two shots of Omicron vaccine, which was not surprising given that the vaccine was matched to the antigen (Figure 4I). However, two shots of Omicron vaccine did not generate higher levels of Omicron-specific antibody responses (Figure 4J). In fact, Omicron-specific antibody responses were similar in both groups at the end of this homologous prime-boost experiment (Figure 4J). To understand how each vaccine affected long-term antibody production, we quantified plasma cells that were specific for the ancestral virus or the Omicron variant (Figure 4K). The results were consistent with the antibody data; the ancestral prime-boost vaccine elicited higher numbers of ancestral-specific plasma cells, and similar number of Omicron-specific plasma cells, relative to the Omicron prime-boost vaccine (Figure 4L-4M). Altogether, these data suggest that in the context of a homologous prime-boost vaccination, Omicron-based vaccines may not necessarily confer superior Omicron-specific immunity, compared to ancestral vaccines.

However, our pre-boost data suggested that with a single prime immunization, an Omicron vaccine may confer a slight advantage over an ancestral vaccine (p=0.06 at pre-boost day 21, Figure 4J). Although this difference was not statistically significant it motivated us to repeat these experiments with a higher number of mice and compare in more detail the primary immune response and protection induced by a single shot of Omicron vaccine (Figure 5A). T cell responses were comparable in both groups that received a single prime (Figure 5B-5C). Similar to our prior experiments, antibody responses tended to be higher when the antigen was matched to its respective vaccine (Figure 5D-5E). This effect was also observed for neutralizing antibody responses (Figure 5F-5G). In particular, the Omicron vaccine prime elicited 19.6-fold improved neutralizing antibody responses against Omicron, relative to the ancestral prime (Figure 5G). Based on these data, we hypothesized that in the context of a single prime immunization, an Omicron vaccine may confer improved protection against Omicron, relative to an ancestral vaccine. To test this hypothesis, we primed K18-hACE2 mice with either an ancestral vaccine or an Omicron vaccine, and at week 2 post-prime, these mice were challenged intranasally with 5×10^4^ PFU of SARS-CoV-2 Omicron variant. Consistent with our prediction, we observed superior protection against Omicron in mice that received a single shot of the Omicron vaccine, relative to mice that received a single shot of the ancestral vaccine (Figure 5H).

**Figure 5.**
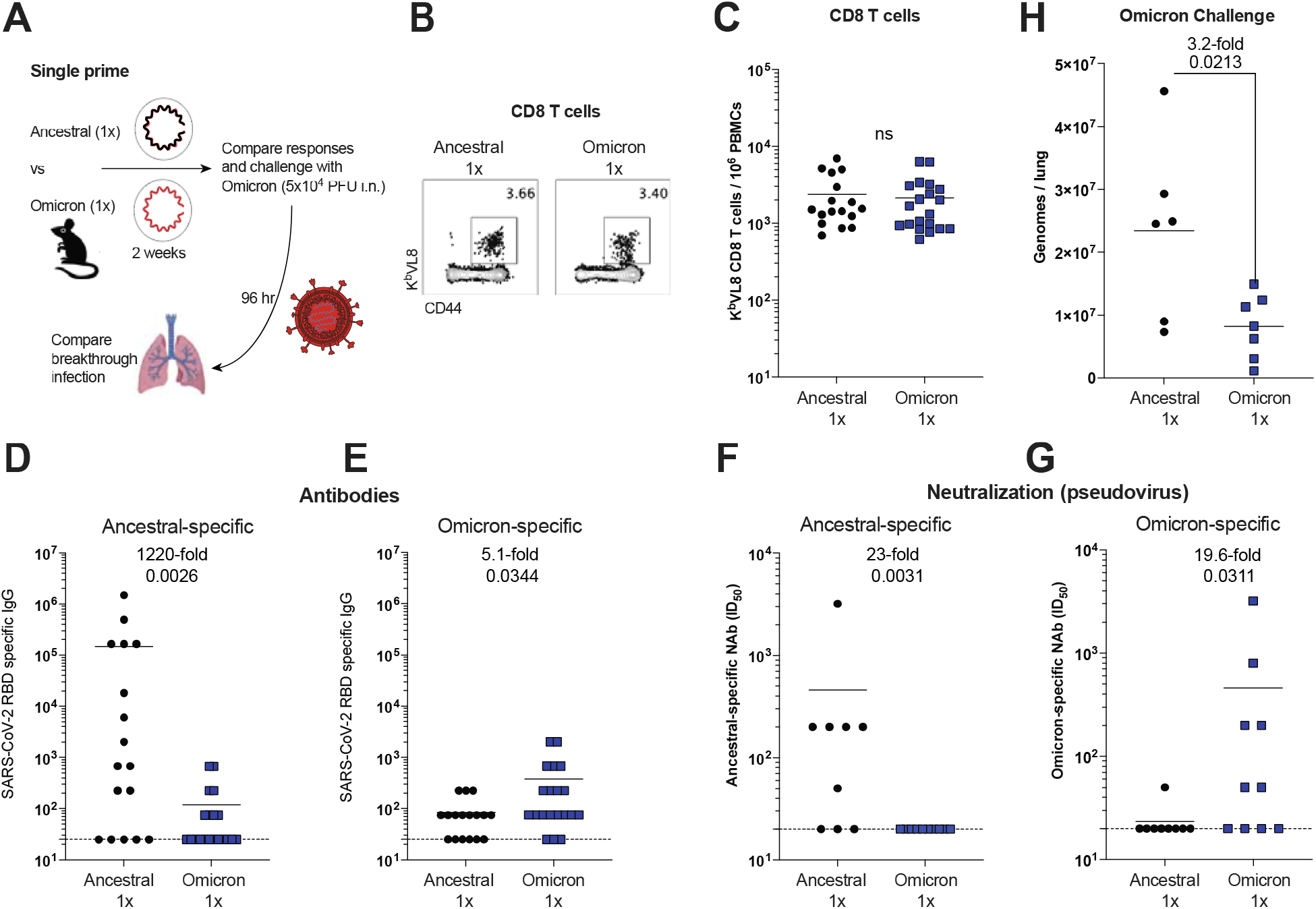
A single prime with an Omicron vaccine results in superior protection against Omicron than a single prime with an ancestral vaccine. (A) Experimental layout. C57BL/6 mice were immunized intramuscularly with 3 μg of an mRNA expressing ancestral spike or Omicron spike. After 2 weeks, immune responses were quantified. (B) Representative FACS plots of SARS-CoV-2 spike–specific CD8 T cells. (C) Summary of SARS-CoV-2 spike–specific CD8 T cells. These CD8 T cells were specific for a conserved CD8 T cell epitope present in both ancestral and Omicron viruses (K^b^VL8). (D) Ancestral spike–specific antibody responses. (E) Omicron spike–specific antibody responses. (F) Ancestral spike–specific neutralizing antibody responses. (G) Omicron spike–specific neutralizing antibody responses. With only a single mRNA prime, there is substantial variability in antibody responses. (H) Viral loads after Omicron challenge. Data from panels A-G used wild type C57BL/6 mice, whereas data from panel H used K18-hACE2 mice (on C57BL/6 background). Two-tailed Mann Whitney test was used. Data from panels C-E are from 3 experiments; the first experiment with 9-10 mice per group, the second experiment with 3-5 mice per group, and the third experiment with 5 mice per group. Data from panels F-G are from 1 experiment with 9-10 mice per group. Data from panel H are from 2 experiments with n=3-4 mice per group (challenges were performed in BSL-3 facilities). All data are shown; dashed lines represent the LOD. Error bars indicate the SEM.

Taken together, the data show that Omicron-specific boosters may not necessarily confer superior immunity relative to ancestral boosters, but in the context of a single prime immunization (in a seronegative host), an Omicron-based vaccine may confer a protective advantage. Notably, the superiority of an Omicron vaccine over an ancestral vaccine critically depends on whether the host is immune to the SARS-CoV-2 spike antigen.

### Pre-existing immunity to the vaccine antigen limits *in situ* antigen expression after mRNA vaccination

So far we have shown that pre-existing immunity can influence responses by mRNA vaccines. We demonstrated that pre-existing antibody can abrogate *de novo* B cell responses so our next aim was to understand a possible mechanism. Historically, the seroprevalence of viral vectors in the human population has hampered their clinical utility, both as vaccines and gene delivery systems. Pre-existing humoral immunity has been especially a limitation for the Ad5 platform, as this virus has infected ∼90% of humans, motivating the development of other vector platforms with lower seroprevalence, such as Ad26 (*10–14*). Specifically, pre-existing antibody generated by prior Ad5 infections limits the amount of antigen that can be expressed after Ad5 immunization (*15*), rendering Ad5 vaccines less immunogenic in seropositive individuals. We interrogated whether a similar phenomenon occurs with mRNA vaccines. In other words, we asked if pre-existing immunity to the antigen encoded by the mRNA vaccine can limit the amount of antigen that is expressed at the site of immunization, following mRNA vaccination.

To answer this simple question, we utilized an mRNA-LNP luciferase system that allowed highly sensitive *in situ* quantification of antigen expression at the site of immunization (the quadriceps muscle) in the presence or absence of pre-existing immunity. We first injected mice with an mRNA-luciferase and after 2 weeks we re-injected these mice with the same mRNA-luciferase to determine whether antigen expression was affected by previous immunization (Figure S4A). As controls, we injected another group of mice with PBS and then with the mRNA-luciferase. Interestingly, mice that received a prior immunization with the mRNA-LNP showed lower antigen expression following re-administration of the same mRNA-LNP in the muscle (Figure S4B-S4C).

To more rigorously evaluate a role for humoral immunity, we performed a passive immunization study (Figure 6A). We primed C57BL/6 mice with mRNA-luciferase, and after 3 weeks, we boosted these same mice with the same mRNA-luciferase to generate a high level of luciferase-specific antibodies. Two weeks after boost, mice were bled and plasma were adoptively transferred into naïve BALB/c mice. As controls, we transferred plasma from mice that were previously immunized with a control mRNA-LNP expressing a different antigen. We included another group of mice that received a different vaccine platform expressing the same antigen (Ad5-luciferase) to rule out potential effects by responses raised against the mRNA-LNP platform itself. A day after adoptive plasma transfer, all mice received an intramuscular injection of mRNA-luciferase and antigen expression was quantified by IVIS. As expected, both mRNA-luciferase and Ad5-luciferase generated luciferase-specific antibody responses in the donor mice (Figure S5). Injection of mRNA-Luciferase into recipient mice resulted in rapid antigen expression by 6 hr post-immunization, and the antigen became undetectable in most mice within a week of immunization (Figure 6B-6C). Interestingly, transfer of luciferase-specific plasma (irrespective of the vaccine platform used) accelerated antigen clearance after mRNA-LNP immunization, especially during the hyperacute phase (6 hr), demonstrating that seropositivity to the vaccine antigen limits the amount of antigen that is expressed at the site of immunization (Figure 6B-6C). These data suggest that pre-existing antibodies can “extinguish” antigen after mRNA vaccination, which may potentially limit the boosting capacity of mRNA vaccines following their re-utilization in the same host. Taken together, we show that pre-existing humoral responses can exert profound effects on the immunogenicity and antigen expression kinetics of mRNA vaccines.

**Figure 6.**
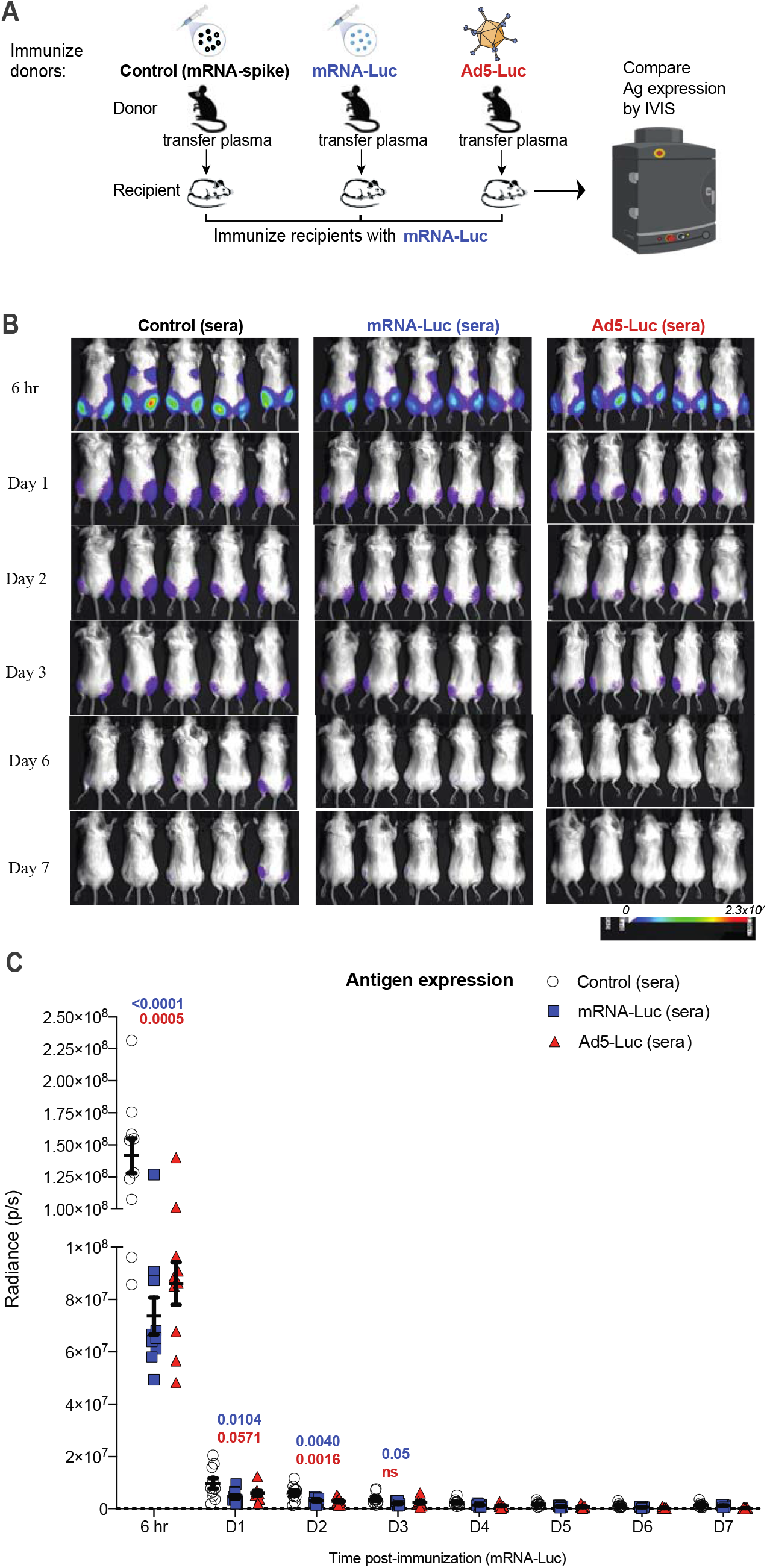
Humoral responses curtail antigen expression following mRNA immunization. (A) Experimental layout. C57BL/6 mice were immunized intramuscularly with 3 μg of an mRNA expressing Luciferase, and after 3 weeks, mice were boosted. We immunized another group of mice with a different vaccine platform expressing the same transgene (Ad5-Luciferase) or with an irrelevant mRNA vaccine expressing a different antigen (mRNA-spike). Two weeks after boost, mice were bled, and plasmas were collected. 400 μL of plasma was adoptively transferred into naïve BALB/c mice via the intraperitoneal route. On the following day, recipient mice received an intramuscular injection of mRNA-luciferase and luciferase expression was quantified by in vivo imaging (IVIS). (B) Bioluminescence images. (C) Summary of transgene expression by IVIS. BALB/c mice (with white coat) were used for improved visualization of luciferase. Data are from one experiment with 10 quadriceps per group (5 mice per group). Experiment was repeated for a total of 2 times, with similar results. Two-way ANOVA test (Dunnett’s multiple comparisons, with adjusted p-value) was used. P-values are indicated in blue (mRNA-control vs mRNA-luciferase), or in red (mRNA-control vs Ad5-luciferase). Error bars indicate the SEM.

## Discussion

mRNA-LNP vaccines have been administered to millions of people worldwide and have shown high efficacy at preventing severe COVID-19. Despite their success, breakthrough infections, variants of concern, and declining antibody levels have warranted the use of boosters, but it remains unclear how pre-existing immunity induced by prior immunization or by infection affects the efficacy of mRNA-LNP vaccines. By analyzing a cohort of individuals who were initially primed with mRNA vaccines, we observed that lower antibody levels before the boost were associated with higher fold-increase in antibody levels after the boost. The same pattern was reported in COVID-19 convalescent patients who subsequently received SARS-CoV-2 vaccines. These data from our human cohorts show that pre-boost antibody titers can be a predictor for antibody responses following vaccination, which has also been shown for Influenza and Herpes Zoster vaccination (*16, 17*).

While the mRNA boost was effective at boosting antibody titers in all individuals, the observation that pre-existing humoral responses downmodulated responses by the mRNA boost suggested a critical mechanistic insight of how B cell responses are regulated following booster vaccination. To further examine this, we performed passive immunization studies in mice, transferring donor-matched plasma from vaccinated humans (before or after vaccination) into mice that were later immunized with mRNA vaccines. Transfer of SARS-CoV-2 spike-immune human plasma into recipient mice abrogated the *de novo* priming of B cell responses following mRNA-spike vaccination. Similar effects were reported in mice that received autologous immune plasma. These data support a model in which antibody “competes” with *de novo* B cell priming. These data advise in favor of extending the prime-boost interval to allow systemic antibody levels to decline, which may facilitate B cell activation after subsequent mRNA vaccination. It has been shown that extended prime-boost intervals elicit more potent antibody responses, but it has been unclear whether this effect is due to intrinsic maturation of immune responses or waning of antibody levels (*15, 18*). Our data bring more clarity to this issue, showing that seropositivity to the vaccine antigen significantly affects the priming of B cell responses. Our data also suggest that the use of monoclonal antibodies or convalescent plasma therapy for COVID-19 may hamper the efficacy of SARS-CoV-2 vaccines, if administered during the time of vaccination. Therefore, our findings may guide clinicians on the optimal time for boosting, ideally after antibody responses contract or after passively transferred antibodies are cleared from circulation, to allow better activation of B cell responses by the mRNA booster.

We also evaluated whether pre-existing antibody affected antigen expression following mRNA vaccination. We performed studies with an mRNA-LNP expressing luciferase, which allowed quantification of antigen expression in the presence or absence of pre-existing immunity. We showed that pre-existing immunity against the antigen encoded by the mRNA vaccine limits antigen expression after mRNA immunization. In the case of adenovirus-based vaccines, pre-existing immunity is known to accelerate the clearance of adenovirus vaccines, limiting the utility of adenoviruses in prime-boost regimens and motivating the use of adenoviruses with lower seroprevalence, such as Ad26. In a seronegative host, adenovirus vaccination results in long-term antigen expression lasting for > 3 weeks; but in a seropositive host, adenovirus vaccination results in transient antigen expression lasting less than a week (*15, 19*). Although both mRNA-based and adenovirus-based vaccines are affected by pre-existing immunity, there are differences in the kinetics of antigen expression. We show that mRNA immunization induces a short-term pulse of antigen, peaking at 6 hr post-immunization and declining to almost undetectable levels within a week of immunization. Pre-existing immunity does not completely prevent the antigen from being expressed after mRNA vaccination, but it could reduce the amount of antigen available to prime the adaptive immune system (Figure 6).

In our studies it is unclear why there were no differences in the magnitude of T cell responses. A possible reason for the discrepancy between B cell and T cell responses is that T cells may require lower amounts of cognate antigen to get primed, and that after a specific “antigen threshold,” more antigen may not necessarily activate more T cells. A prior clinical study in volunteers who received dose-escalating doses of the Pfizer vaccine (BNT162b1) showed a pattern consistent with this notion; antibody responses tend to be higher with higher mRNA doses, but the same dose-dependent pattern is not observed for T cell responses (*20*). Future studies will evaluate in more detail the antigen level requirements for B cells and T cells, and determine why immune plasma abrogated B cell responses but not T cell responses.

Due to the growing interest in booster vaccines, our studies also evaluated how pre-existing immunity affects the immunogenicity of booster vaccines based on the ancestral virus or the Omicron variant. It is still controversial whether Omicron-based vaccines would be necessary, and there are ongoing vaccine trials evaluating this proof of concept. Our data bring more clarity to this issue, as we show that boosting with an Omicron vaccine may not necessarily confer an immunological advantage over boosting with an ancestral vaccine, consistent with recent studies (*1–3, 21*). Like some of those prior studies, our studies suggest original antigenic sin as a possible mechanism for the limited capacity of Omicron boosters to elicit neutralizing antibody. However, we now show that in the context of a single prime, an Omicron vaccine may confer a slight protective advantage against Omicron relative to an ancestral vaccine. This relative advantage of an Omicron vaccine is lost in a seropositive host, suggesting that the relative superiority of an Omicron vaccine depends on whether the host has been exposed to SARS-CoV-2. Thus, it is reasonable to hypothesize that in a seronegative population that may be exposed to Omicron, an Omicron vaccine may confer improved protection relative to an ancestral vaccine.

We also evaluated immune responses after homologous prime-boost immunizations comparing an ancestral vaccine versus an Omicron-based vaccine. Priming and boosting with the same ancestral vaccine elicited higher levels of ancestral-specific antibody responses, relative to priming and boosting with the same Omicron vaccine. This is an expected outcome, since the vaccine antigen is matched. However, an unexpected finding is that priming and boosting with the Omicron vaccine did not elicit higher levels of Omicron-specific responses, relative to priming and boosting with the ancestral vaccine. This effect could be due to the low structural stability of the Omicron spike protein, which has been reported to be a poor immunogen (*22*). Our data show that the Omicron spike protein is effective at priming Omicron-specific B cell responses, but following a booster, more structurally stable spike proteins may be more effective at expanding pre-existing B cell responses. As variant-specific vaccines are being considered worldwide, it is important to better understand potential benefits and drawbacks of variant-specific vaccines, including their structural stability and immunogenicity *in vivo*. A limitation of developing variant-specific vaccines for use as boosters is that SARS-CoV-2 is a continuously evolving virus, so it may be difficult to keep up with every future variant of concern (VOC). Another limitation is that the spike protein of certain VOCs, including Omicron, may not be optimal immunogens, which may hinder their ability to generate appropriate neutralizing antibody responses (*22*). The development of the current SARS-CoV-2 vaccines was in part possible by introducing stabilizing amino acids in the ancestral spike protein (two prolines at position 986-987), demonstrating that structural stability can sometimes be more important than exact antigenic matching (*23, 24*).

Taken together, there are three critical points from this study. First, pre-existing humoral responses can limit B cell responses following mRNA vaccination. This observation suggests that following vaccination or infection it may be beneficial to wait until antibody levels contract before administering an mRNA boost. Similarly, in the context of monoclonal antibody therapy or convalescent plasma therapy, it would be beneficial to wait until antibody is cleared before administering an mRNA vaccine. Second, pre-existing immunity to the antigen encoded by the mRNA vaccine accelerates the clearance of the mRNA vaccine at the site of immunization, limiting the amount of antigen available to prime to adaptive immune system. This insight may help to devise strategies to extend the duration of gene expression by mRNA-LNPs used in gene therapy, for example by transiently downmodulating immune responses with immunosuppressive drugs to facilitate the serial re-utilization of the mRNA-LNPs. Third, we show that boosting with an Omicron vaccine does not confer an immunological advantage relative to boosting with ancestral vaccine. However, in the context of a single prime, an Omicron vaccine confers a relative immunological advantage. These data are important for understanding how pre-existing immunity modulates responses to mRNA-LNPs, offering insights to improve their efficacy.

## Resource availability

### Lead contacts

Further information and requests for resources and reagents should be directed to and will be fulfilled by the lead contact Pablo Penaloza-MacMaster.

### Materials availability

For mRNA vaccine access, contact Pablo Penaloza-MacMaster.

### Data and code availability

This article does not report original code. The article includes all the data sets and analyses generated for this study. Any additional information required to reanalyze the data reported in this paper is available from the lead contact upon request.

### Experimental models and subject details

#### Animals and ethics statement

Mice were purchased from Jackson laboratories and were housed at Northwestern University or University of Illinois at Chicago (UIC) animal facility. All procedures were performed with the approval of the center for comparative medicine at Northwestern University and the UIC IACUC. Adult mice, approximately half females and half males, were used for the immunogenicity experiments included in this study. For the challenge studies, female mice were used.

In the human studies, all protocols used for participant recruitment, enrollment, blood collection, sample processing, and immunological assays with human samples were approved by the IRB of Northwestern University (STU00212583). All individuals voluntarily enrolled in the study by signing an informed consent form after receiving detailed information about the study.

### Method Details

#### Mice and vaccinations

6-8-week-old K18-hACE2 mice were used for challenge studies, as done previously (*25*). These mice express human ACE2 on the keratin 18 promoter. These mice were purchased from Jackson laboratories (Stock No: 034860). Mice were immunized intramuscularly (50 μL per quadriceps) of mRNA-LNPs encoding the respective SARS-CoV-2 spike protein diluted in sterile PBS, at 3 μg per mouse, as done previously (*25, 26*). All other experiments were performed with 6-8-week-old wild type mice from Jackson laboratories (C57BL/6, Stock No: 000664; or BALB/c, Stock No: 000651).

#### SARS-CoV-2 virus and infections

SARS-Related Coronavirus 2, Omicron Isolate was obtained through BEI Resources, NIAID, NIH: SARS-Related Coronavirus 2, Isolate hCoV-19/USA/MD-HP20874/2021 (Lineage B.1.1.529; Omicron Variant), NR-56461, contributed by Andrew S. Pekosz. This virus was propagated and tittered on Vero-E6 cells (ATCC). Vero-E6 cells were passaged in DMEM with 10% Fetal bovine serum (FBS) and Glutamax. Cells less than 20 passages were used for all studies. Virus stocks were expanded in Vero-E6 cells following a low MOI (0.01) inoculation and harvested after 96 hr. Titers were determined by plaque assay on Vero-E6 cell monolayers. Viral stocks were used after a single expansion (passage = 1) to prevent genetic drift. K18-hACE2 mice were anesthetized with isoflurane and challenged with 5×10^4^ PFU of virus intranasally. Mouse infectious were performed at the University of Illinois at Chicago (UIC) following BL3 guidelines with approval by the UIC Institutional Animal Care and Use Committee (IACUC).

#### SARS-CoV-2 quantification in lungs

Lungs were isolated from infected mice and homogenized in sterile PBS. RNA was isolated with the Zymo 96-well RNA isolation kit (Catalog #: R1052) following the manufacturer’s protocol. SARS-CoV-2 viral burden was measured by RT-qPCR using Taqman primer and probe sets from IDT with the following sequences: Forward 5’ GAC CCC AAA ATC AGC GAA AT 3’, Reverse 5’ TCT GGT TAC TGC CAG TTG AAT CTG 3’, Probe 5’ ACC CCG CAT TAC GTT TGG TGG ACC 3’. A SARS-CoV-2 copy number control was obtained from BEI (NR-52358) and used to quantify SARS-CoV-2 genomes.

#### Reagents, flow cytometry and equipment

Single cell suspensions were obtained from PBMCs or tissues. Dead cells were gated out using Live/Dead fixable dead cell stain (Invitrogen). MHC class I monomers (K^b^VL8, VNFNFNGL) were used for detecting virus-specific CD8 T cells, and were obtained from the NIH tetramer facility located at Emory University. MHC monomers were tetramerized in-house. The VNFNFNGL epitope is located in position 539-546 of the SARS-CoV-2 spike protein. Cells were stained with fluorescently-labeled antibodies against CD8α (53-6.7 on PerCP-Cy5.5), CD44 (IM7 on Pacific Blue), K^b^VL8 (APC). Fluorescently labeled antibodies were purchased from BD Pharmingen, except for anti-CD44 (which was from Biolegend). Flow cytometry samples were acquired with a Becton Dickinson Canto II or an LSRII and analyzed using FlowJo v10 (Treestar).

#### SARS-CoV-2 and luciferase spike-specific ELISA

Binding antibody titers were quantified using ELISA as described previously (*15, 27, 28*), using the respective proteins as coating antigen. Briefly, 96-well, flat-bottom MaxiSorp plates (Thermo Fisher Scientific) were coated with 1 μg/mL of the respective protein for 48 hours at 4°C. Plates were washed 3 times with wash buffer (PBS plus 0.05% Tween 20). Blocking was performed with blocking solution (200 μl PBS plus 0.05% Tween 20 plus 2% BSA) for 4 hr at room temperature. 6 μl of plasma samples were added to 144 μl of blocking solution in the first column of the plate, 3-fold serial dilutions were prepared for each sample, and plates were incubated for 1 hr at room temperature. Plates were washed 3 times with wash buffer. To determine mouse allotype-specific antibody, different primary antibodies were used; biotin anti-mouse IgG1^[a]^ specific antibody (BD-Pharmingen, MN 553500) and biotin anti-mouse IgG1^[b]^ specific antibody (BD-Pharmingen, MN 553533), which were diluted 1:1000 in blocking solution and were then added to the plates and incubated for 1 hr at room temperature. Plates were washed 3 times with wash buffer and added streptavidin-HRP (Southern Biotech, 7105-05) diluted 1:400 in blocking buffer and incubated for 1 hr at room temperature. After washing plates 3 times with wash buffer, 100 μl/well SureBlue Substrate (SeraCare) was added for 1 min. The reaction was stopped using 100 μl/well KPL TMB Stop Solution (SeraCare). Absorbance was measured at 450 nm using a Spectramax Plus 384 (Molecular Devices). Ancestral SARS-CoV-2 spike used in ELISAs was produced in-house using a plasmid produced under HHSN272201400008C and obtained from BEI Resources, NIAID, NIH: vector pCAGGS containing the SARS-related coronavirus 2; Wuhan-Hu-1 spike glycoprotein gene (soluble, stabilized); NR-52394. The ancestral receptor binding domain (RBD) protein used as coating antigen was made in-house; produced under HHSN272201400008C and obtained through BEI Resources (NIAID, NIH: Vector pCAGGS Containing the SARS-Related Coronavirus 2, Wuhan-Hu-1 Spike Glycoprotein Receptor Binding Domain (RBD), NR-52309). The Omicron RBD protein used as coating antigen was purchased from RayBiotech. The luciferase protein used as coating antigen was purchased from Sigma (L9420).

### mRNA-LNP vaccines

We synthesized mRNA vaccines encoding for the codon-optimized SARS-CoV-2 spike protein from USA-WA1/2020 or Omicron. Constructs were purchased from Integrated DNA Technologies (IDT) or Genscript, respectively, and contained a T7 promoter site for in vitro transcription of mRNA. The sequences of the 5′- and 3′-UTRs were identical to those used in a previous publication (*26*). All mRNAs were encapsulated into lipid nanoparticles using the NanoAssemblr Benchtop system (Precision NanoSystems). mRNA was dissolved in Formulation Buffer (catalog NWW0043, Precision NanoSystems) and then run through a laminar flow cartridge with GenVoy ILM (catalog NWW0041, Precision NanoSystems) encapsulation lipids at a flow ratio of 3:1 (RNA: GenVoy-ILM), with a total flow rate of 12 mL/min, to produce mRNA–lipid nanoparticles (mRNA-LNPs). mRNA-LNPs were evaluated for encapsulation efficiency and mRNA concentration using RiboGreen assay using the Quant-iT RiboGreen RNA Assay Kit (catalog R11490, Invitrogen, Thermo Fisher Scientific). mRNA to express luciferase was purchased from TriLink Biotechnologies (CleanCap® FLuc mRNA, L-7602, CleanCap®Firefly Luciferase) and was encapsulated into lipid nanoparticles using the aforementioned protocol.

### In vivo bioluminescence

Mice were imaged at various times after immunization. To quantify luciferase expression, luciferin (GoldBio, catalog no. LUCK-100; weight of 10 μl/g) was administered intraperitoneally 15 minutes before imaging. Mice were anesthetized and imaged for 2 minutes using a KINO IVIS Imager (Spectral Instruments Imaging). Region of interest (ROI) bioluminescence was used to quantify response. Each leg (quadriceps) was treated as a an individual site of immunization (or individual replicate).

### Pseudovirus neutralization assays

SARS-CoV-2 pseudovirus neutralization assays were performed as described previously (*15, 29*). In brief, we utilized a SARS-CoV-2 spike pseudotyped lentivirus kit was obtained through BEI Resources, NIAID, NIH (SARS-Related Coronavirus 2, Wuhan-Hu-1 Spike-Pseudotyped Lentiviral Kit V2, NR-53816). We used HEK-293T– hACE2 cells as targets (BEI Resources, NIAID, NIH, NR-52511). Serial plasma dilutions were incubated with the SARS-CoV-2 spike pseudotyped lentivirus. Cells were then lysed using luciferase cell culture lysis buffer (Promega). Luciferase reaction was performed using 30 μl of cell lysis buffer (Promega). The reaction was added to 96-well black optiplates (PerkinElmer) and luminescence was quantified using a PerkinElmer Victor3 luminometer.

### B cell ELISPOT

Antibody secreting cells (ACS) were enumerated similar to a prior paper, but using spike protein as coating antigen instead of viral lysate (*27*). In brief, SARS-CoV2 spike-specific ASC were quantitated by ELISPOT using 96-well Multiscreen filter plate (MSHAN4B50, Millipore Ireland BV). Plates were coated with 5 μg/mL of the SARS-CoV-2 spike protein and incubated overnight at 4°C. After incubation, plates were washed once with PBS containing 0.05% Tween-20 (PBS-T) and twice with PBS. Plates were blocked by incubating plates with RPMI containing 10% fetal bovine serum (FBS) for 2 hr at room temperature. Single suspensions of splenocytes at 60×10^6^ cells/mL were prepared in medium (RPMI supplemented with 10% FBS, 1% penicillin/streptomycin, and 1% L-glutamine and 0.05 mM of B-Mercaptoethanol). After incubation, blocking medium was replaced with 100 μl/well of fresh medium and added 50 μl of single cell suspension to the first row, and serially diluted 3-fold down the plate. Plates were incubated for 8 hr at 37°C in a 5% CO_2_ incubator. Plates were washed with PBS and PBS-T, and incubated with 100 μl of biotinylated anti-mouse IgG antibody diluted 1:1000 in PBS-T with 1% FBS for two days at 4°C. Antibody was removed by flicking plates followed by washing four times with PBS-T, and 100 μl of horseradish peroxidase (HRP) conjugated avidin D (A-1004, Vector Laboratories, Burlingame, CA) was added per well at 5 μg/mL in PBS-T with 1% FCS and incubated for 1 hr at room temperature. Plates were washed three times with PBS-T and PBS before adding 100 μl of freshly prepared chromogen substrate. The substrate was prepared by adding 0.15 ml of AEC solution (3-amino-9-ethyl carbazole, MP Biochemical#195039) at a concentration of 20 mg/mL in dimethylformamide (Sigma, St. Louis, MO) to 10 ml of 0.1M sodium acetate buffer (pH = 4.8). This solution was filtered through a 0.2 μm membrane and 100 μl of 3% H_2_O_2_ was added immediately before use. Plates were incubated with substrate for 8 min until spots appeared and the reaction was stopped by rinsing plates with water. Plates were allowed to dry and spots were counted.

#### Quantification and statistical analysis

Statistical analyses are indicated on the figure legend. Dashed lines in data figures represent limit of detection. Statistical significance was established at p ≤0.05 and the statistical test used is indicated otherwise in figure legends. Data were analyzed using Prism (Graphpad).

## Author Contributions

T.D. and S.S. performed the immunogenicity experiments. J.R. performed the SARS-CoV-2 challenge experiments. L.V. and I.K. recruited human volunteers and harvested plasma samples for analyses. P.P.M. designed the experiments and secured funding. P.P.M. wrote the paper with feedback from all authors.

## Acknowledgements

We thank Dr. Andreas Wieland for discussions. This work was possible with a grant from the National Institute of Biomedical Imaging and Bioengineering (NIBIB U54 EB027049) and a grant from the National Institute on Drug Abuse (NIDA, DP2DA051912) to P.P.M.

## Supplemental Figures

**Figure S1.**
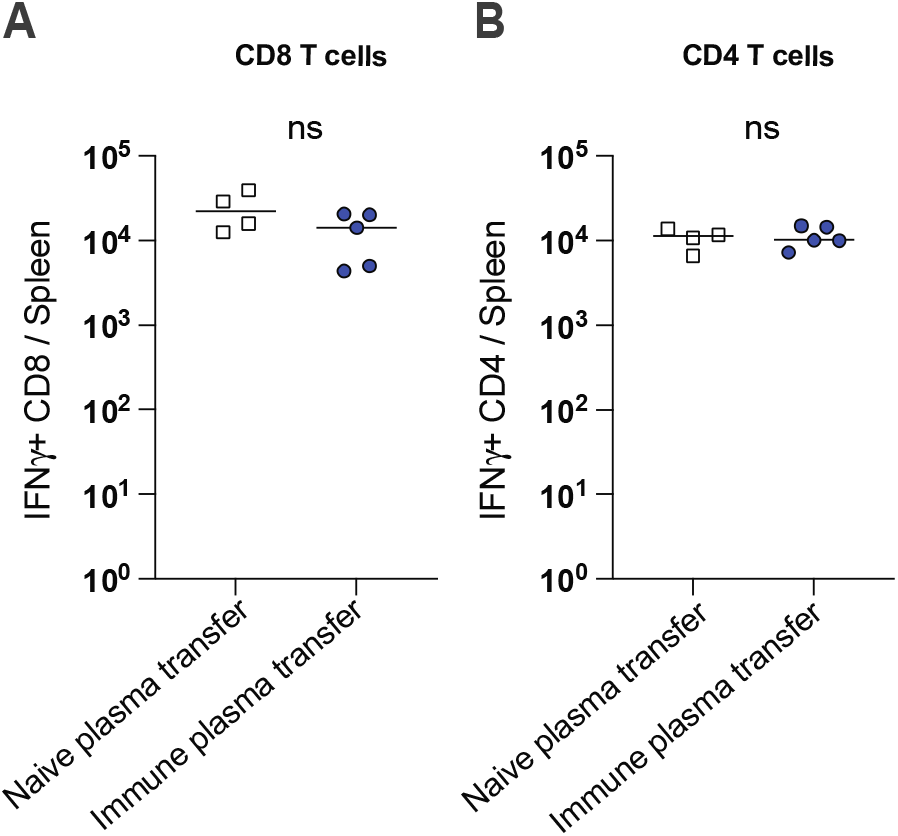
Plasma from vaccinated mice does not abrogate T cell responses. Plasmas were harvested from C57BL/6 mice that were vaccinated with mRNA-spike (two doses). 400 μL of plasma were adoptively transferred via the intraperitoneal route into BALB/c mice. On the following day, all mice were immunized intramuscularly with 3 μg of an mRNA expressing SARS-CoV-2 spike; and T cell responses were quantified by intracellular cytokine staining (ICS) at day 21 post-vaccination. Naïve plasmas were used as controls. Splenocytes were incubated with overlapping SARS-CoV-2 peptide pools for 5 hr at 37°C in the presence of GolgiStop and GolgiPlug to detect SARS-CoV-2–specific CD8+ T cell responses (A) and CD4+ T cell responses (B). Two-tailed Mann Whitney test was used. Data are from an experiment with 4 mice that received naïve plasma and 5 mice that received post-vaccine plasma (week 2 post-boost); experiment was repeated for a total of 2 times, with similar results; dashed lines represent the LOD. Error bars indicate the SEM.

**Figure S2.**
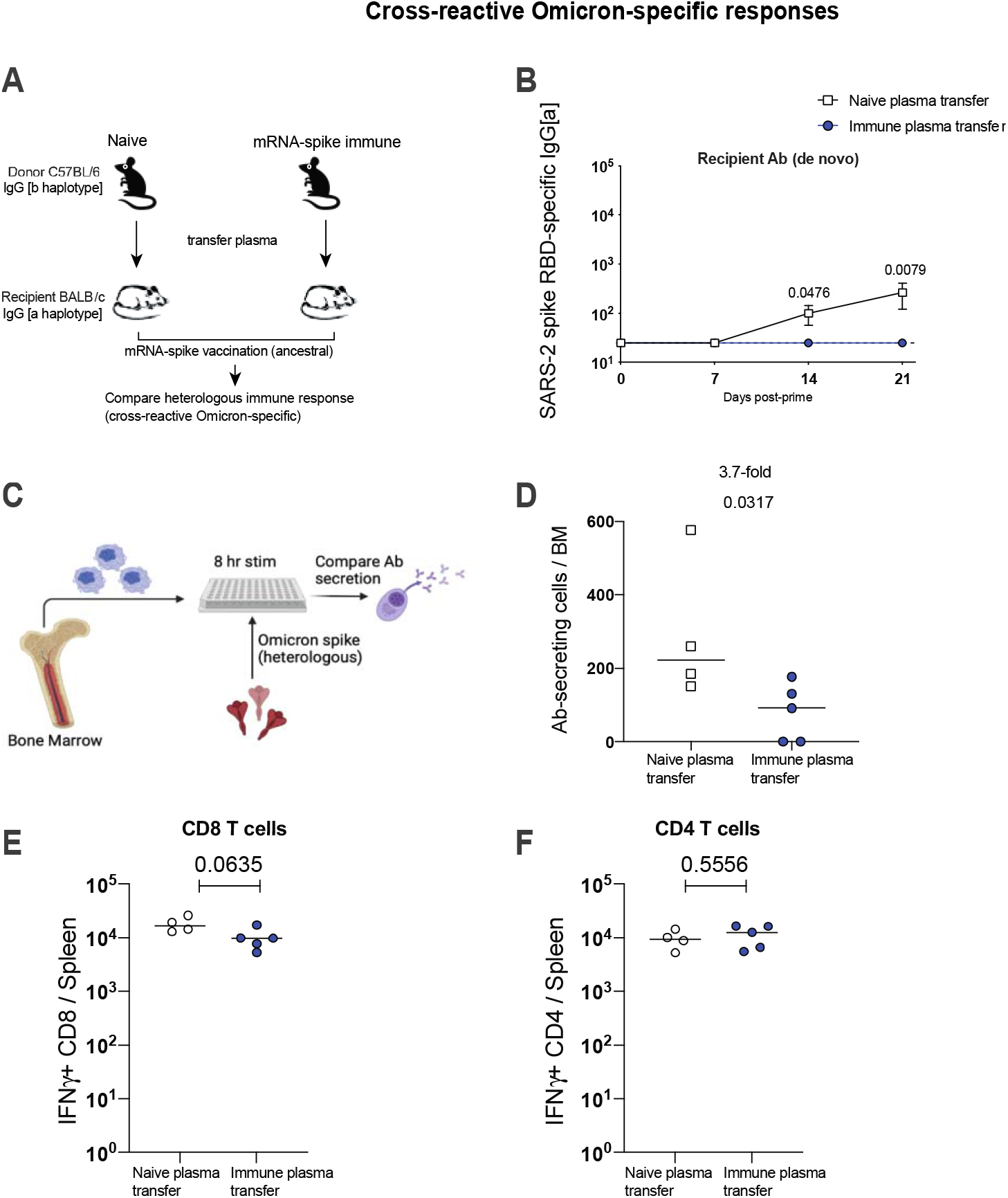
Plasma from vaccinated mice abrogates cross-reactive (Omicron-specific) antibody responses. Plasmas were harvested from C57BL/6 mice that were vaccinated with mRNA-spike (two doses). 400 μL of plasma was adoptively transferred via the intraperitoneal route into BALB/c mice. On the following day, all mice were immunized intramuscularly with 3 μg of an mRNA expressing SARS-CoV-2 spike (ancestral sequence); and cross-reactive Omicron-specific immune responses were quantified. Naïve plasmas were used as controls. (B) Recipient-derived Omicron-specific antibody. (C) Experimental layout for detection of antibody secreting cells specific for ancestral or Omicron spike. (D) Omicron-specific antibody secreting cells in bone marrow. (E-F) T cell responses were quantified by intracellular cytokine staining (ICS) at day 21 post-vaccination. Splenocytes were incubated with overlapping SARS-CoV-2 peptide pools (derived from the Omicron variant) for 5 hr at 37°C in the presence of GolgiStop and GolgiPlug to detect SARS-CoV-2–specific CD8+ T cell responses (E) and CD4+ T cell responses (F). Two-tailed Mann Whitney test was used. Data are from an experiment with 4 mice that received naïve plasma and 5 mice that received post-vaccine plasma (week 2 post-boost); experiment was repeated for a total of 2 times, with similar results; dashed lines represent the LOD. Error bars indicate the SEM.

**Figure S3.**
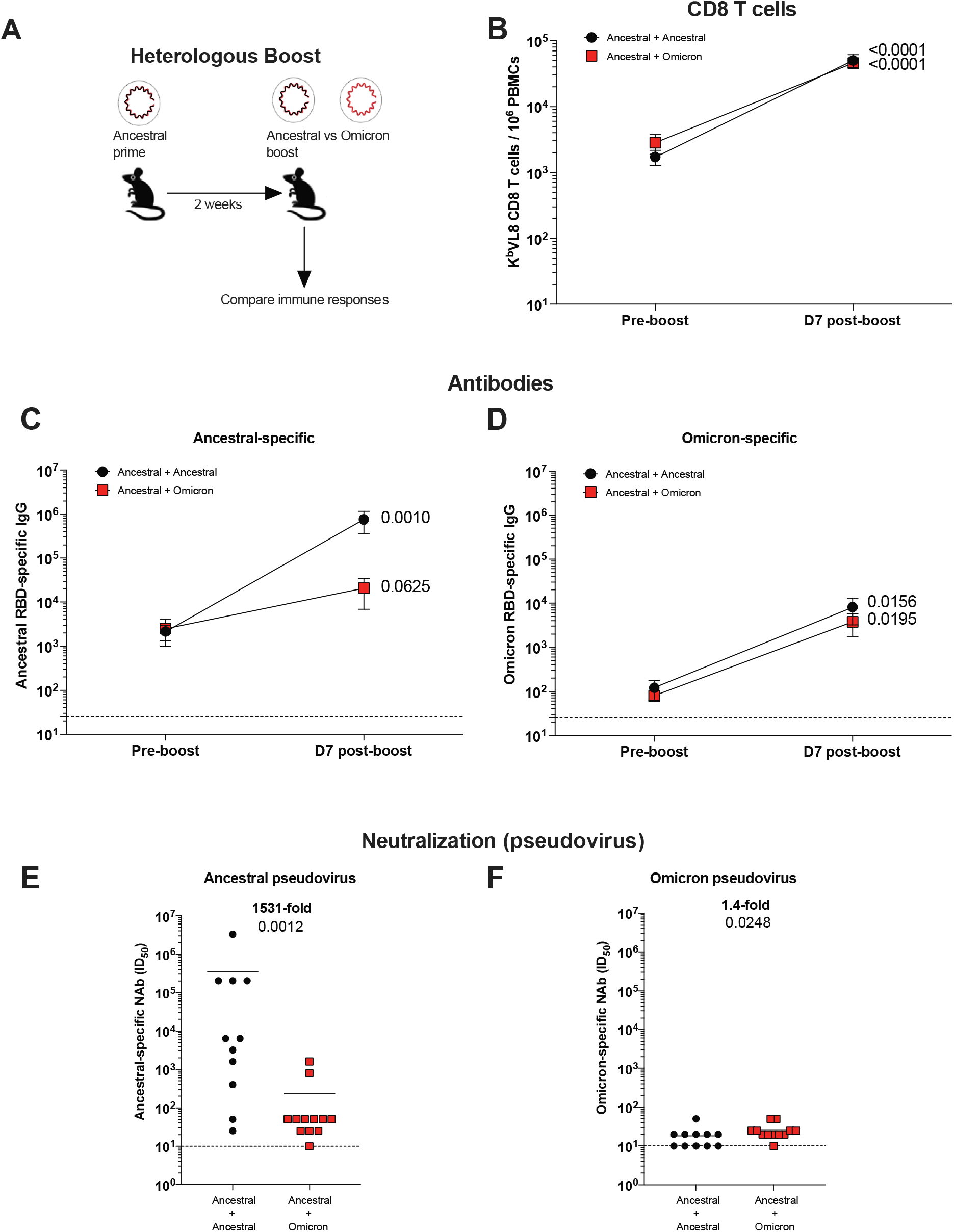
A heterologous Omicron boost is not superior to a homologous ancestral vaccine boost. (A) Experimental layout. C57BL/6 mice were immunized intramuscularly with 3 μg of an mRNA expressing ancestral spike. After 3 weeks, mice were boosted with either an mRNA expressing ancestral spike, or an mRNA expressing Omicron spike. Immune responses were quantified. (B) Summary of SARS-CoV-2 spike–specific CD8 T cells. These CD8 T cells were specific for a conserved CD8 T cell epitope present in both ancestral and Omicron viruses (K^b^VL8). (C) Ancestral spike–specific antibody responses. (D) Omicron spike–specific antibody responses. (E) Ancestral spike–specific neutralizing antibody responses. (F) Omicron spike–specific neutralizing antibody responses. Wilcoxon matched-pairs signed rank test was used for panels B-D, comparing pre-boost and post-boost responses for each vaccine regimen. For all other plots, two-tailed Mann Whitney test was used. Data are combined from two experiments; one experiment with 6-7 mice per group and another experiment with 5 mice per group. Dashed lines represent the LOD (in one experiment the LOD was 25). Error bars indicate the SEM.

**Figure S4.**
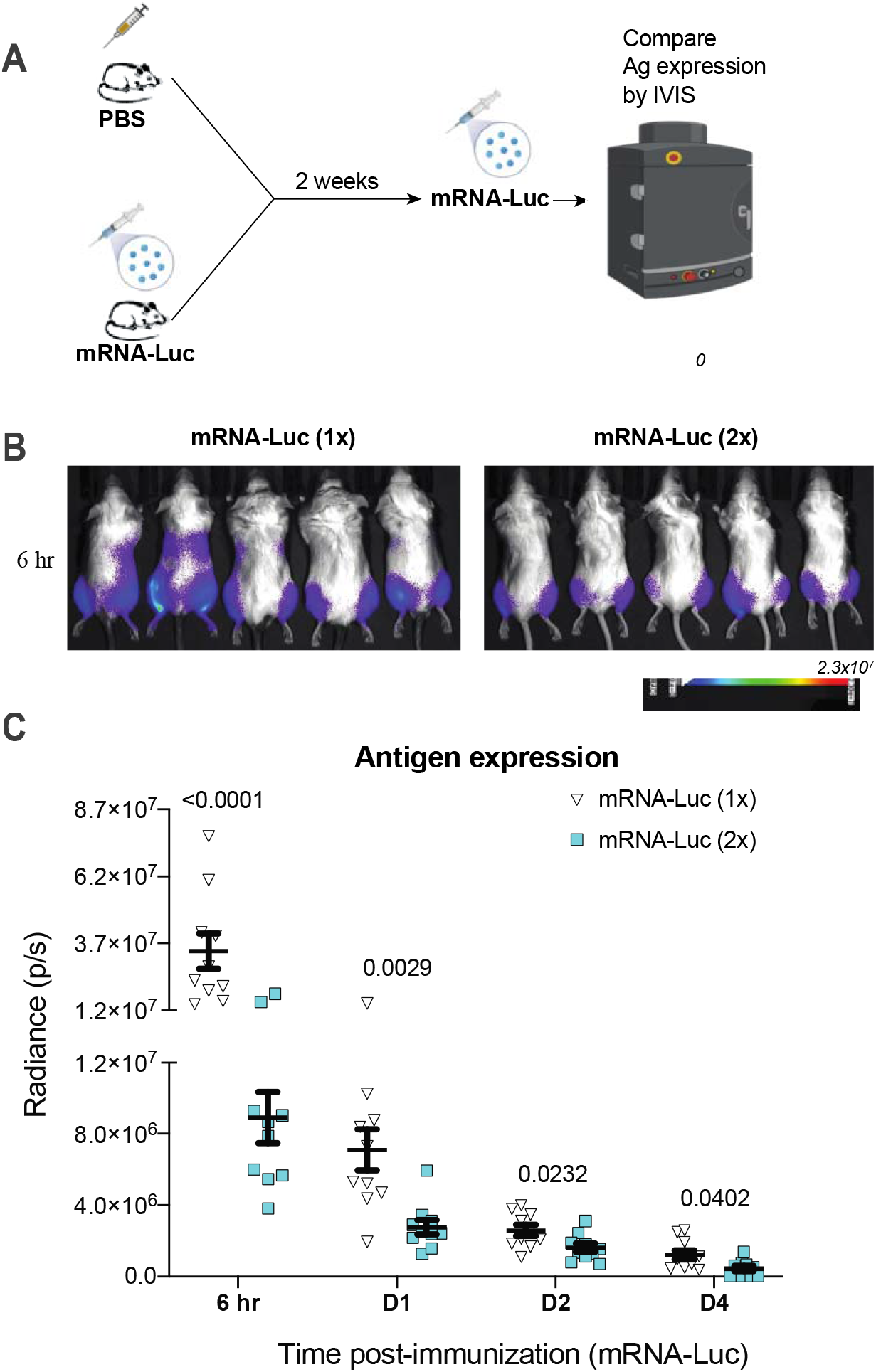
Boosting with mRNA results in lower antigen expression, relative to priming with mRNA. (A) Experimental layout. BALB/c mice were immunized intramuscularly with 3 μg of an mRNA expressing Luciferase (mRNA-Luc). After 2 weeks, mice were boosted with the same mRNA, and luciferase expression was quantified by IVIS. (B) Bioluminescence images at 6 hr. (C) Summary of transgene expression by in vivo bioluminescence imaging. BALB/c mice (with white coat) were used for improved visualization of luciferase. Two-tailed Mann Whitney test was used. Data are from one experiment with 10 quadriceps per group (5 mice per group). Experiment was repeated for a total of 2 times, with similar results. Error bars indicate the SEM.

**Figure S5.**
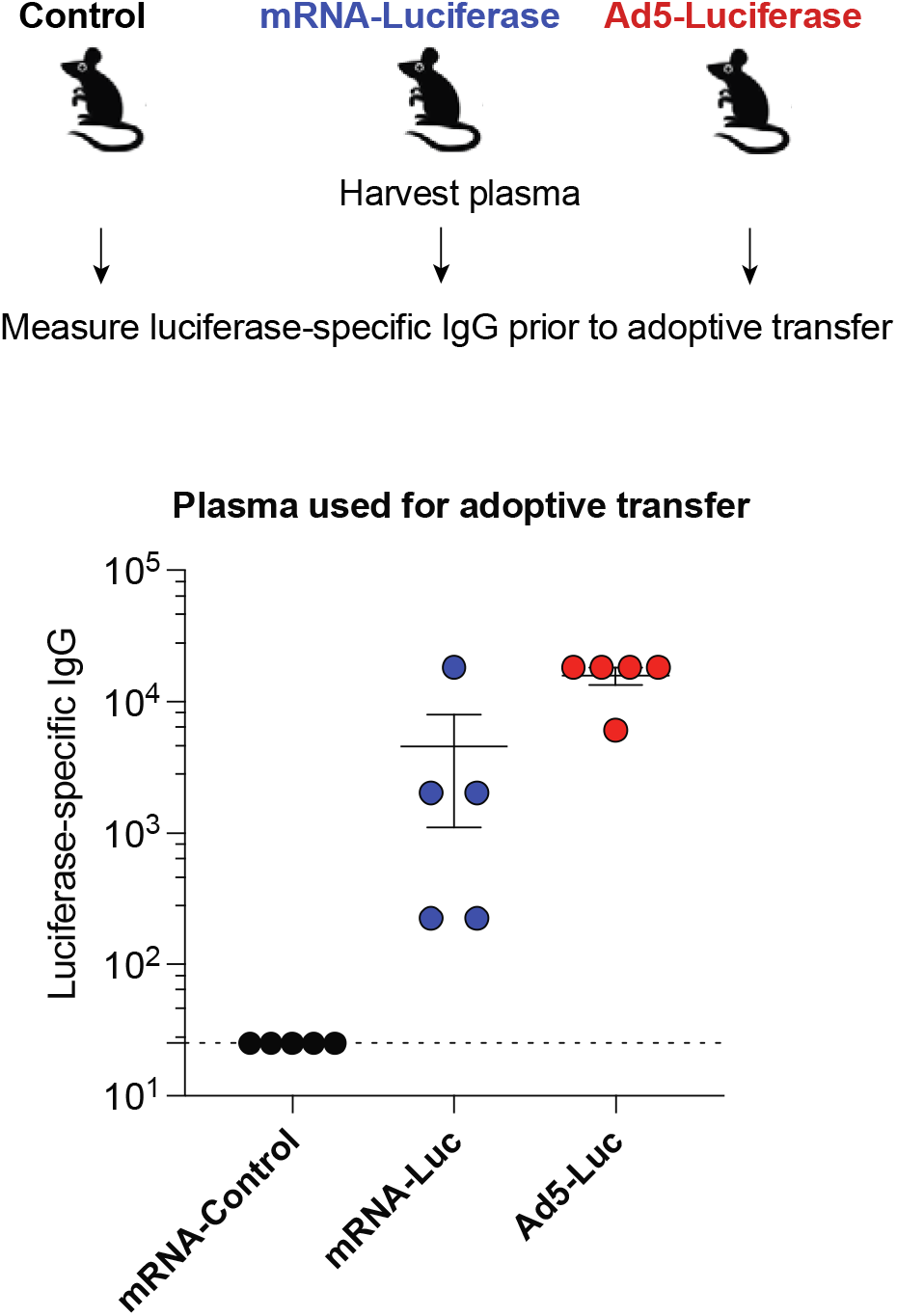
mRNA-Luciferase immunization induces luciferase-specific antibody responses. C57BL/6 mice were immunized intramuscularly with 3 μg of an mRNA expressing Luciferase, and after 3 weeks, mice were boosted. We immunized another group of mice with a different vaccine platform expressing the same transgene (Ad5-Luciferase) or with an irrelevant control mRNA vaccine expressing a different antigen (mRNA-spike). Two weeks after boost, mice were bled, and luciferase-specific antibodies were quantified in plasma. These plasma samples were used in adoptive transfers in Figure 6. Luciferase-specific antibody responses are shown. Data are from one experiment with 5 donor mice. Experiment was repeated for a total of 2 times, with similar results. Error bars indicate the SEM.

